# Spatial and temporal contexts drive visual temporal integration via spatiotemporal normalisation

**DOI:** 10.64898/2025.12.26.696562

**Authors:** Ljubica Jovanovic, Pascal Mamassian

**Author notes:** Corresponding Author: Ljubica Jovanovic.

## Abstract

The visual system is known to integrate sensory information across both space and time, but these dimensions are often studied separately. Here we provide novel behavioural evidence for an early integration over space and time, and propose a computational model of this integration that is biologically plausible. We designed a controlled task that forces human participants to integrate information over both space and time to perform well. Stimuli were temporally segmented rings of oriented elements and participants had to detect a missing element in the ring. We found that temporal integration was enhanced when successive visual events were temporally proximate, spatially aligned to form a continuous contour, but also, and more surprisingly, when they were brief. Notably, the spatial effects on integration are temporally asymmetric: integration is facilitated when more events are presented earlier in the sequence, and even more so when they form a collinear contour. These findings indicate that temporal integration is sensitive to both the spatial configuration and the number of visual events presented over time, supporting a continuous rather than discrete process at early visual stages. All of these results are well accounted for by a computational model in which temporal integration arises from the overlap of internal signals generated by individual visual events. These signals are shaped by a spatially and temporally tuned divisive normalisation mechanism and integrated via coincidence detection. Predictions from this model match human performance in seminal past work as well as novel stimulus displays, offering an account of how spatio-temporal interactions determine what we perceive and when.

## Introduction

The human visual system is bombarded with a continuous flow of sensory signals. In order to extract relevant information and drive appropriate behaviour, these signals are integrated across space and time, thereby reducing noise at the cost of some precision loss. The integration over space and time shares some common computational principles^1–6^, but traditionally these two dimensions have been studied separately^2^. Here we leveraged human psychophysics and computational modelling to offer a unifying account of these integrations. We revisited a precise psychophysical paradigm^7,8^ that was designed to measure the temporal integration of the visual system and extended it to characterise both the spatial and the temporal integrations and their interactions. These findings led us to develop a computational model of spatio-temporal integration of visual information that is based on divisive normalisation, a canonical neural computation^9–12^. The model describes generic computations of the early visual system that affects all subsequent visual processes, and ultimately contributes to what we perceive and when we perceive it.

Integrating sensory signals over temporal windows helps to consolidate the relevant information before a perceptual decision is made^13–16^. While these integration windows have been posited more than a century ago, we are still missing a description of the mechanism that determine which information is included and how it is integrated^17–20^. A promising starting point is found at the neural level in humans, where temporal receptive fields have been studied using various model-based approaches across different sensory modalities^21–24^. In particular, it is now clear that the sensory signals undergo different nonlinear transformations, and that these non-linearities have a functional role^22,23,25^. In the human visual cortex, these non-linearities have been modelled as a Delayed Normalisation and explain how successive stimuli are combined without saturating the system^23,25^. The appeal of the model lies in the fact that it is an instance of divisive normalisation, a canonical neural computation widely observed across the brain and in different sensory systems^10–12,26^. This framework proposes that the initial response of a neuron is divided by the summed activity of a pool of neurons that includes itself and surrounding ones. By modelling the normalisation over time and delaying the effects of the normalisation pool, the Delayed Normalisation model parsimoniously accounts for effects of stimulus duration, contrast and repetition on the complex nonlinear neural dynamics^25^. While this work provides important insights into how information is integrated over time at different levels of sensory information processing, past research has disregarded how space contributes to the temporal integration for simplicity.

The interaction of spatial and temporal contexts significantly increases the complexity of processing of natural visual scenes. Let us take the example of contour integration that groups isolated elements into perceptually coherent shapes^27–29^. It is supported by both lateral connections in primary visual cortex (V1) and feedback from higher visual areas^30–39^. Collinearity of elements promotes their grouping^40^, and can increase sensitivity to weak signals along the contour^41,42^. From previous work, we know that temporal properties influence spatial integration, for instance temporal synchrony favours collinearity detection and grouping over space^43–48^. However, less is known about how contour integration based on collinearity affects temporal integration^49,50^, limiting our understanding of how the brain integrates both space and time to interpret dynamic and complex visual scenes.

Here we first provide novel evidence for inseparability of spatial and temporal integration in the human visual system. Using a temporally segmented ring (TSR) stimulus, we establish an intricate relationship between contour collinearity and temporal integration. Collinearity in our displays improves temporal integration, but this effect is modulated by how information is made available over time, even at very short time scales (∼ 50 ms). These new behavioural data help us develop a computational model that describes putative computations underlying spatio-temporal integration of visual information. We extend the Delayed Normalisation model^23,25^ with a spatial weighting and feature tuning of the normalisation pool^51^ to account for responses of the visual system to our TSR displays. Integration over time of these responses is implemented by quantifying the temporal overlap between predicted signals^52–54^. The model offers a unifying account of key features of both seminal results in temporal and spatial integration studies and our novel findings using a spatio-temporal display. Finally, we directly compare the model’s predictions to human performance for a novel set of stimuli, establishing the generalisability of the model and offering a novel tool to study how visual percepts build up over time.

## Results

To investigate the dynamics of spatial and temporal context on temporal integration, we performed a controlled psychophysical experiment while manipulating different properties of a specifically-tailored stimulus. The stimulus, that we call the ”temporally segmented ring” (TSR), consists of a ring of oriented elements that are split in two groups. Each group contains one half of the elements, and the two groups are presented in rapid succession to probe temporal integration. When one element is removed from one of the groups, detection of the missing element is easy if the temporal gap between the two groups is small, and hard if it is large. In our experiments, participants were presented with two such TSR stimuli, one standard where all the oriented elements were present and the other one a test where one element was missing. They were then asked to identify the interval with a missing element (sequential 2AFC paradigm, **Fig.1a**). We manipulated the temporal gap between the two groups of a TSR stimulus and the duration of the first group (see Methods for details).

**Figure 1.**
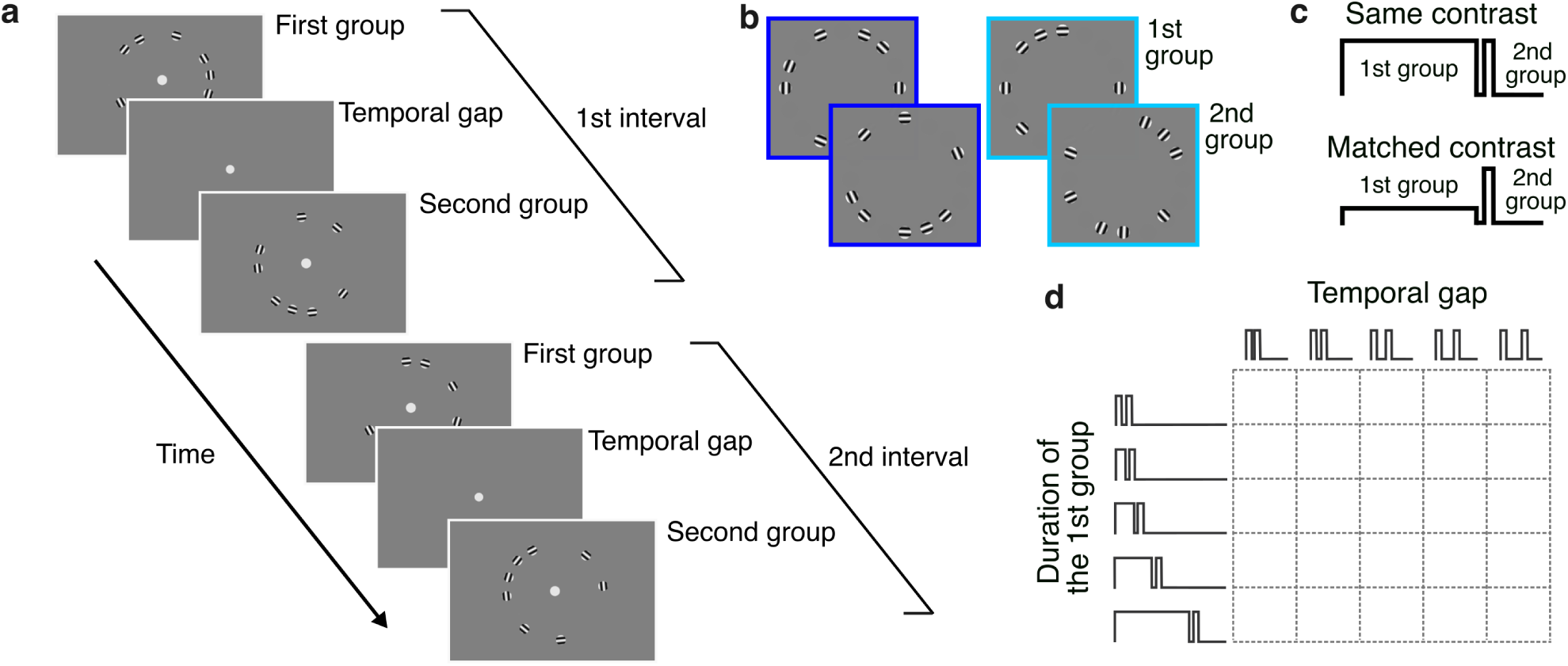
Procedure in Experiment 1. **a**. Stimuli were temporally segmented rings (TSR) consisting of an array of grating elements arranged on a virtual circle around the fixation point. The gratings were randomly split in two equal-size groups, presented briefly one after the other, separated by a temporal gap. On each trial, two TSR stimuli were presented in two consecutive intervals. In a two alternative forced-choice paradigm (2AFC), one of the two intervals contained a test TSR where one grating element was missing, and participants had to identify the interval that contained this test TSR. **b**. Two spatial conditions were tested. The orientation of the gratings in the second group was either tangential to the virtual circle, forming a continuous contour (dark blue), or orthogonal to the virtual circle, forming a disjoined contour (light blue). **c**. In two different sessions, two contrast conditions were tested. The contrast of the two groups was either identical (100%), or the contrast of the first group was adjusted individually to match the perceived contrast of the second group (decreased contrast for longer durations of the first group). **d**. Duration of the first group (10 − 160 ms) and temporal gap between the two groups (0 − 40 ms) were systematically varied in each block.

### Temporal integration is modulated by fast spatial grouping

As the temporal gap between the two groups of gratings is increased, we anticipate that the detectability of the ring with the missing element will get harder. In a first Experiment, we varied temporal gap between the two groups (0–40 ms), and also independently the duration of the first group of gratings (10–160 ms; **Fig.1d**). Intuitively, we might expect that performance should decrease as temporal gap increases, and increase as duration increases. This latter prediction results from the fact that as duration increases, the first group of gratings becomes more visible. As expected, temporal integration performance dropped with the duration of the temporal gap (GLMM, type III Wald *χ*^2^(1) = 1650, *p <* 0.01; **Fig.2**). More surprisingly however, performance decreased with the duration of the first group (*χ*^2^(2) = 1077, *p <* 0.01). This result is actually in line with some previous work^7^, and suggests that the first group of elements can inhibit the second group rather than the two being simply added. This property will be critical when we build our model in a later section. In addition, the effect of the duration was smaller at longer temporal gaps (*χ*^2^(2) = 13072, *p <* 0.01).

**Figure 2.**
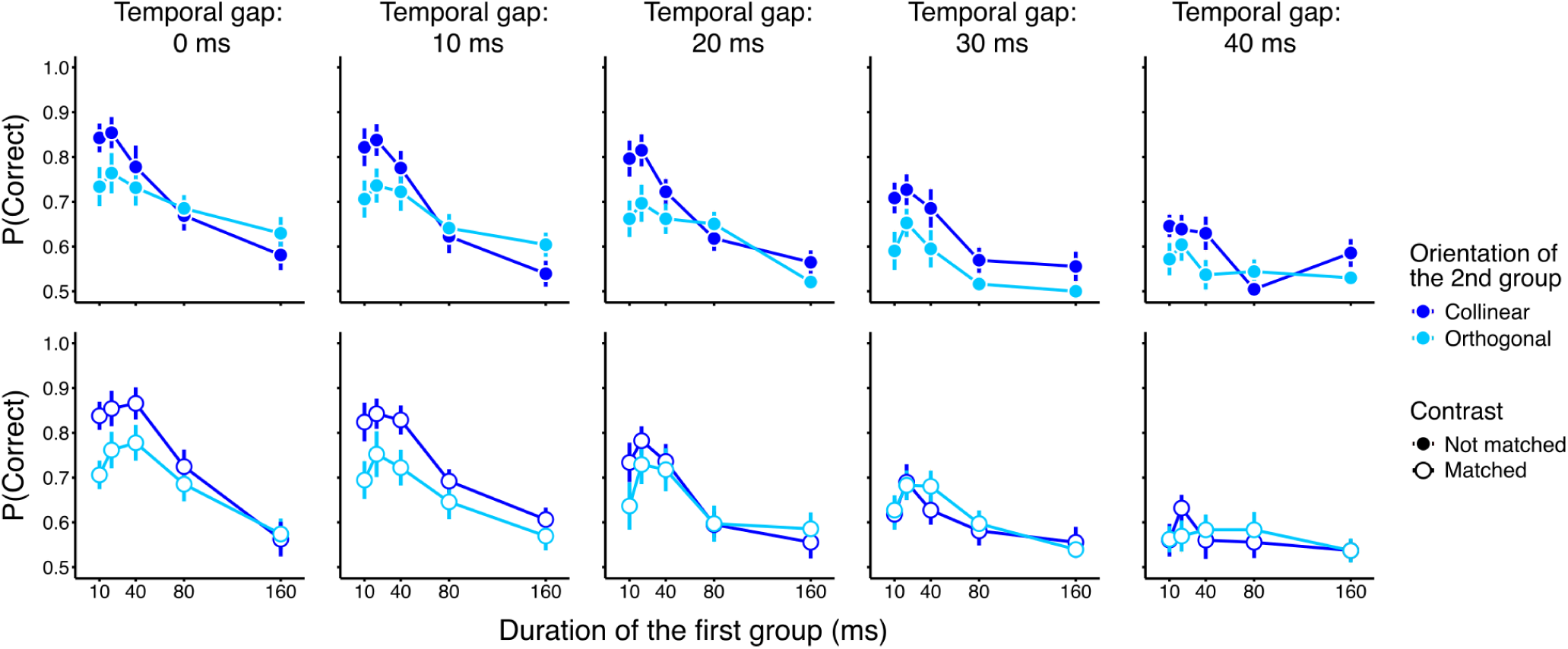
Results of Experiment 1. The fraction of correct detection of the interval containing the missing element is plotted as a function of the duration of the first group of elements of a TSR stimulus. Each panel correspond to one set of conditions. The columns represent the five temporal gap conditions, the rows the effect of contrast of the first group, and the colour of the symbols whether the second group of elements were aligned or orthogonal to the contour. Performance decreases as a function of temporal gap, and, more surprisingly, as a function of the duration of the first group. Performance was better when all the gratings were collinear to the contour, indicating an interaction of spatial and temporal contexts. Finally, matching the perceived contrast between the two groups of elements improved slightly performance for longer durations of the first group. Error bars are standard errors of the mean.

**Figure 3.**
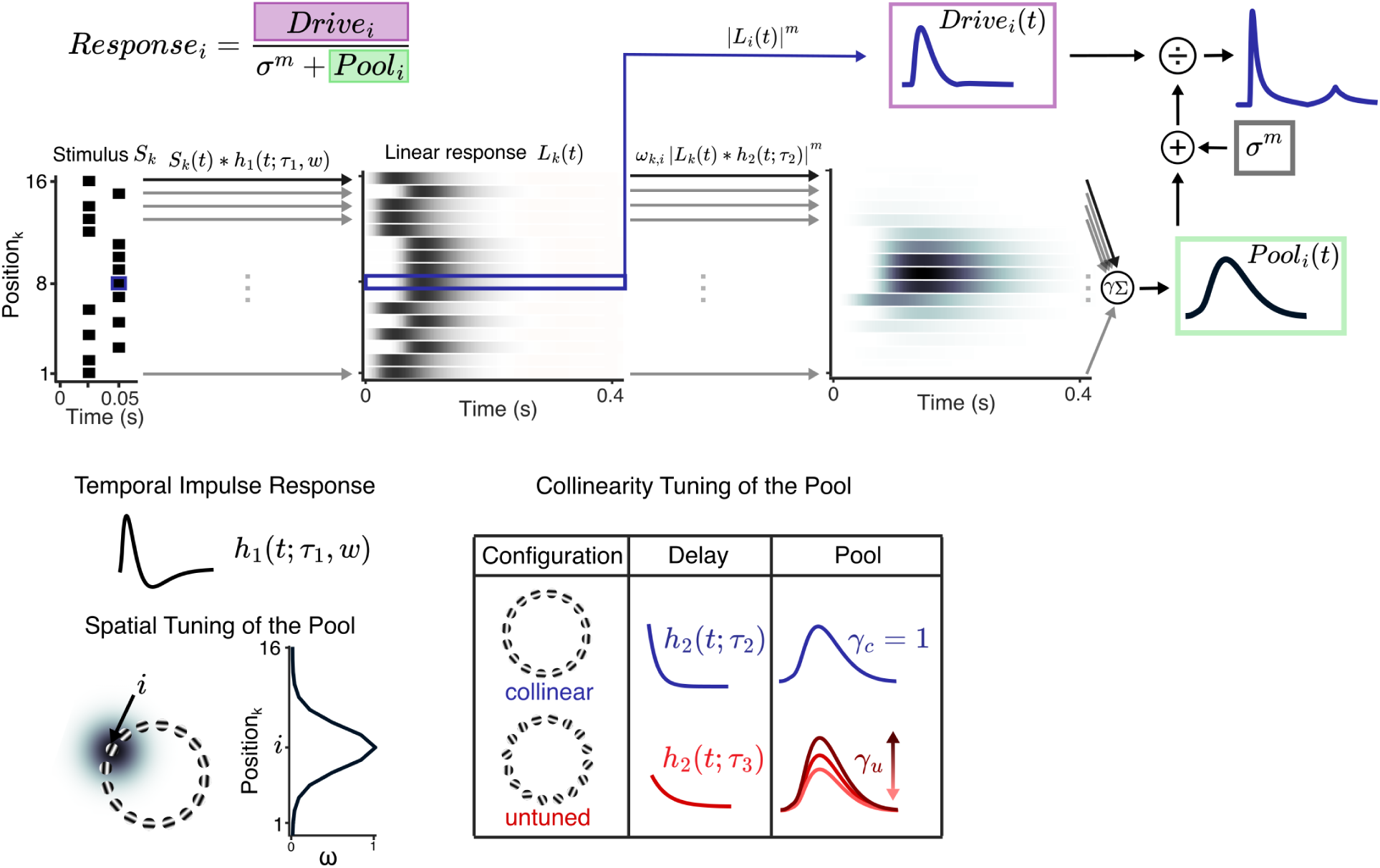
Illustration of the Tuned Delayed Normalisation Model. The temporal part of the model is the Delayed Normalisation^10,23,25^, that has been extended here to include spatial properties of the visual stimulus in the normalisation pool. The illustration depicts the model’s predicted response for one element i along the ring (here the 8*^th^* element, shown in dark blue), that was presented for 10 ms in the second group, in the experimental condition where all the gratings were collinear. The response’s drive is the convolution of the stimulus time-course with a temporal impulse response (*h*_1_, difference of two gamma functions), rectified and exponentiated to the power *m* (purple rectangle). A normalised response is then obtained by dividing the drive by the sum of a semi-saturation constant (*σ*) and the normalisation pool (green rectangle), both raised to the same power *m*. The normalisation pool is modelled as a sum of spatially weighted delayed linear responses to all the gratings presented in the display. Spatial weight (*ω*) is computed as a 2D Gaussian distribution (bottom left of the figure), centered on the grating, whose size is determined by standard deviation *σs*. The delay is achieved by a convolution of the linear response to the gratings with an exponentially decaying function *h*_2_. Elements in the ring have different time constants of the exponential function depending on their orientation (*τ*_2_ for collinear shown in blue, and *τ* 3 for other stimuli shown in red), and contribute differently to the pool (*γ_c_* fixed to 1 and *γ_u_* a free parameter; shown in table).

In this first experiment, we also compared two spatial conditions. The gratings’ orientations were either always aligned with the virtual circle (forming a collinear contour when grouped), or aligned in the first group and orthogonal in the other (forming an disjoined contour; **Fig.1b**). Importantly, this spatial manipulation was crossed with all the temporal conditions described above, thereby allowing us to test the interaction of temporal and spatial contexts on temporal integration. The results revealed a strong effect of the spatial context on the temporal integration. The detectability of a missing element along the contour was overall better when the orientations of the gratings in both groups were aligned with the virtual circle (*χ*^2^(1) = 273.5, *p <* 0.01). Spatial and temporal contexts also interacted: both the effects of the duration of the first group (*χ*^2^(2) = 405.8, *p <* 0.01) and of the temporal gap (*χ*^2^(1) = 273.81, *p <* 0.01) were smaller when the second group contained orthogonal rather than collinear elements (**Fig**.**2**, light vs dark blue symbols).

For brief stimulus presentations, the duration of the stimulus is related to its perceived contrast^15^. To determine whether the effects of durations reported above are related to physical or perceived contrast, the experiment was actually divided in two separate sessions (see Methods). In one session, the contrast of the first group of a TSR stimulus was fixed to the same value as that for the second group, and in another session, it was adjusted as a function of the group’s duration to match the perceived contrast of the second group (individually adjusted in a preliminary calibration experiment, **Fig**.**1c**). The effect of this adjustment on temporal integration depended on the temporal context, as reflected in significant interactions between the type of stimuli (adjusted or not) and the duration of the first group and the temporal gap. When the contrast of stimuli was matched rather than constant (open symbols in **Fig**.**2**), the performance declined more slowly with duration of the first group (*χ*^2^(2) = 759.3, *p <* 0.01), and longer temporal gaps (*χ*^2^(1) = 79.91, *p <* 0.01). We also found higher-order interactions, suggestive of a complex relationship between the spatial and temporal context and contrast of the first group (see Supplementary Information). In summary, using a temporally segmented ring stimulus, we found some clear effect of the spatial configuration of the ring on the temporal integration of its elements. This interaction of spatial and temporal attributes of the stimulus indicates a fast processing of spatial properties that occurs concomitantly with temporal processing. In the next section, we develop a computational model that combines space and time to explain the complexity of the spatio-temporal interactions that we found experimentally. Our model accounts for the effect of the temporal gap between the two groups of elements of the stimulus, and more critically for the detrimental effect of increasing the duration of the first group. The model also makes quantitative predictions that reproduce the effects of our contrast manipulations.

### Temporal integration with a Tuned Delayed Normalisation model

The computational model predicts visual responses to sequentially presented stimuli and quantifies the probability of their integration. Specifically, to predict visual responses to the displays, we appropriated the Delayed Normalisation model^10,23,25^ and extended it to include an enriched normalisation pool. The response to each element of the ring is modulated by the activity resulting from all the elements, with a contribution that varies in strength depending on their spatial and temporal proximity, as well as their orientation. Temporal integration is modelled as a coincidence detection across these normalised responses.

### Tuned Delayed Normalisation

The Delayed Normalisation model^23,25^ predicts the neural response time course by applying a sequence of operations on the visual signal that consists of linear filtering, exponentiation and normalisation.

The time course of each element *i* (individual grating) of the stimulus is filtered by a temporal impulse response

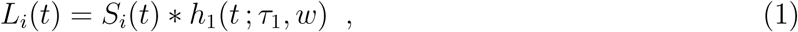

where *S_i_* denotes the time course of the contrast change for the element *i*, and *h*_1_ is the temporal impulse response, defined as a weighted difference of two gamma functions

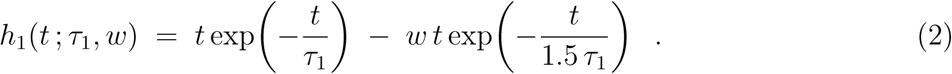

The time constant *τ*_1_ determines the shape of the response and *w* is the weight of the second gamma function.

The drive of an element *i* is the rectified and exponentiated linear response to that element

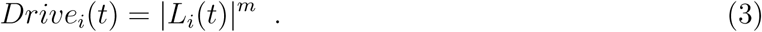

The responses are normalised by the sum of the activity of the normalisation pool and a semi-saturation constant. In previous work, the pool has been modelled as a delayed version of the linear response, rectified and exponentiated. The delay is implemented as a convolution of the linear response with an exponential decay function *h*_2_

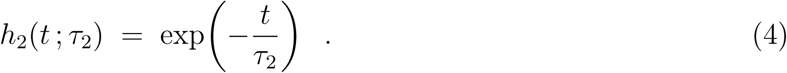

We enriched the normalisation pool to account for spatial interactions and the relative orientation between gratings. The spatial interactions are modelled as a weighting across space, parameterised as a two-dimensional Gaussian function, centered to the grating’s location. The weight of the response to a nearby grating *k* contributing to the group of the *i^th^* grating is thus

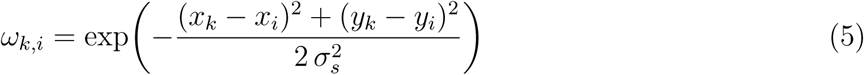

where *x_k_* and *y_k_* are the spatial coordinates of the center of the element *k*. The mean of the 2D Gaussian function is the center of the element being normalised (*x_i_* and *y_i_*), and we assume it to be circular symmetric (same variance *σ*^2^ in *x* and *y* and covariance 0).

To account for the effect of orientation of temporal integration, we introduce parameters allowing for different contributions of the neighbouring units to the normalisation pool. Specifically, the responses of the neighbouring units contribute differently to the pool according to their orientation relative to the virtual circle along which they are positioned. We allow the time constant of the spatial effect to be different for collinear and non-collinear elements (*τ*_2_ and *τ*_3_ respectively)

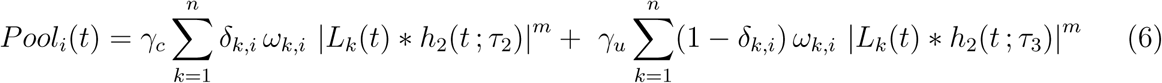

where *γ_c_* and *γ_u_* are weights controlling the strength of the effects of collinear and non-collinear elements, respectively. The Kronecker delta *δ_k,i_* determines how the response to the element at position *k* contributes to the pool of element *i*, as follows

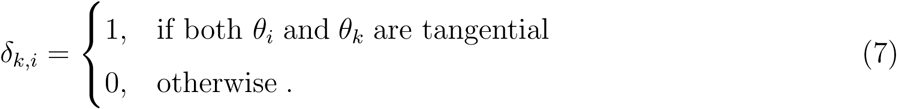

The normalised response (*NR*) for each element is then simply its drive divided by the sum of the normalisation pool and a normalisation constant *σ_c_*

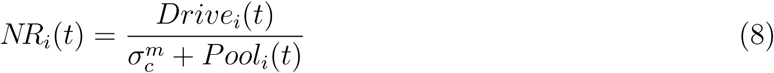

### Temporal integration

To describe how the distributed time-varying activity is integrated, we modelled temporal integration as coincidence detection. Each individual normalised response is thresholded (logistic function Φ, with parameters *p*_1_ and *p*_2_), to model the time course of its detection as a function of response strength

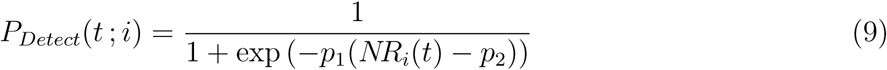

Then, these probabilities are multiplied to obtain the probability of their co-occurrence over time, and then summed across the stimulus duration

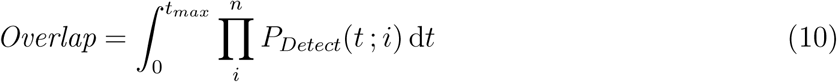

The summed probability of co-occurrence over time corresponds to the duration of the temporal overlap of the detected signal, and it is considered as evidence of temporal integration. In other words, the longer the elements are perceived together, the greater the probability of correctly detecting if there is a missing element. The predicted performance is obtained by passing the integrated temporal overlap through a logistic function Φ, with parameters *p*_3_ and *p*_4_ and *g* = 0.5 (controlling for the lower asymptote).

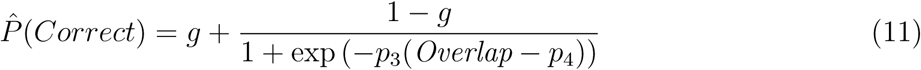

The Delayed Normalisation model for the stimulus where all the elements are aligned has five parameters. These parameters control the temporal impulse response shape (difference of two gamma functions, *τ*_1_ and *w*), an exponent (*m*), a normalisation constant (*σ_c_*) and a time constant of an exponential function (*τ*_2_). All parameters except the normalisation constant were fixed to values previously reported in the literature to provide the best fit for responses of the human visual cortex (for more details see Methods). We allowed for the normalisation constant (*σ_c_*) to vary as a free parameter, as it controls the contribution of the normalisation to the overall response, and might depend on the specific visual displays or task demands. To account for spatial tuning and collinearity effects, we included three free parameters, corresponding to the standard deviation of the spatial tuning of the normalisation (*σ_s_*), a possibly different delay for the non-collinear elements (*τ*_3_), and relative different strengths of normalisation between collinear and untuned pool (*γ_u_*; the other weight *γ_c_* is fixed to 1).

In addition to the parameters used for normalisation, to model performance in the temporal integration task, we included two free parameters to control the thresholding operation (logistic function, *p*_1_, *p*_2_), and two more parameters of the logistic function (*p*_3_, *p*_4_) to transform the temporal overlap into the probability of choosing the correct interval.

### Modelling the data

The model replicates the main features of the human behaviour obtained in Experiment 1 (**Fig. 5**). In particular, the probability of correctly detecting a missing element decreases with an increase of duration of the first group or an increase of the temporal gap, and there is an interaction between these two effects. Performance improves when gratings from both groups are collinear, and contrast-matched displays (lower contrast of the first group) leads to better performance for longer durations of the first group when duration of the first group and temporal gap are short. Intuitively, the model performs poorly when the duration of the first group is long because in that case the delayed normalisation pool is still very strong by the time the second group of elements is displayed, thereby reducing the visibility of both the first and the second group. Spatially, the temporal dynamics of the neural response to each element are modulated by the activity of adjacent elements, such that increased normalisation strength leads to a shortened response duration or a delayed onset and peak. Furthermore, the improvement of performance for collinear displays comes from two parameters: a stronger weight of the activity of orthogonal gratings to the normalisation pool (*γ_u_ > γ_c_*) and a faster normalisation by collinear elements, suppressing the activity of the collinear gratings earlier (*τ*_2_ *> τ*_3_, see Table 1). The spatial normalisation and feature tuning properties of the model constitute significant evolutions of the delayed normalisation models^23,25^. To test whether additional complexity is justified given our experimental results, we fit the data with a reduced model, and perform a nested hypothesis testing. We keep the structure of the decisional part of the model (detection and integration) identical, and modify the normalisation mechanism, following the model structure reported in previous work^25^. Specifically, response to each grating is normalised only by a delayed version of that response, and with equal intensity and time-constant regardless of its relative orientation. A nested hypothesis testing (likelihood ratio test) indicates that the full model describes the data significantly better (*λ_Nested_* = −1496.51, *λ_F ull_* = −1490.8, *χ*^2^(3) = 11.42, *p <* 0.01). Even though the model can qualitatively predict the effects of the duration of the first group, temporal gap and contrast, it fails to account for the differences in the relative orientations of the gratings across the two groups (see Supplementary Information).

**Figure 4.**
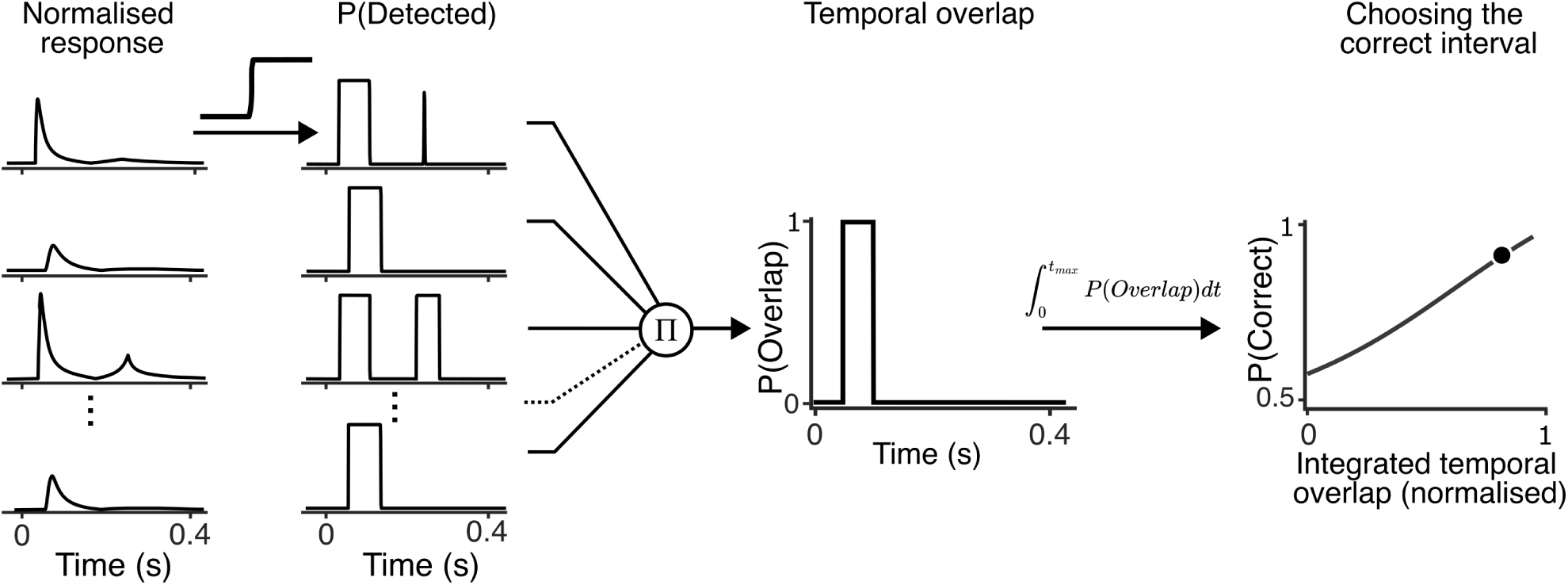
Illustration of temporal integration. Predicted normalised response for each grating of the display (first column) is thresholded, to model probability of its detection over time (*P_Detect_*, second column). These responses are multiplied, to obtain probability of their concurrence over time (*P_Overlap_*, third column). Finally, temporal overlap integrated over time determines probability of detection of the missing element.

**Figure 5.**
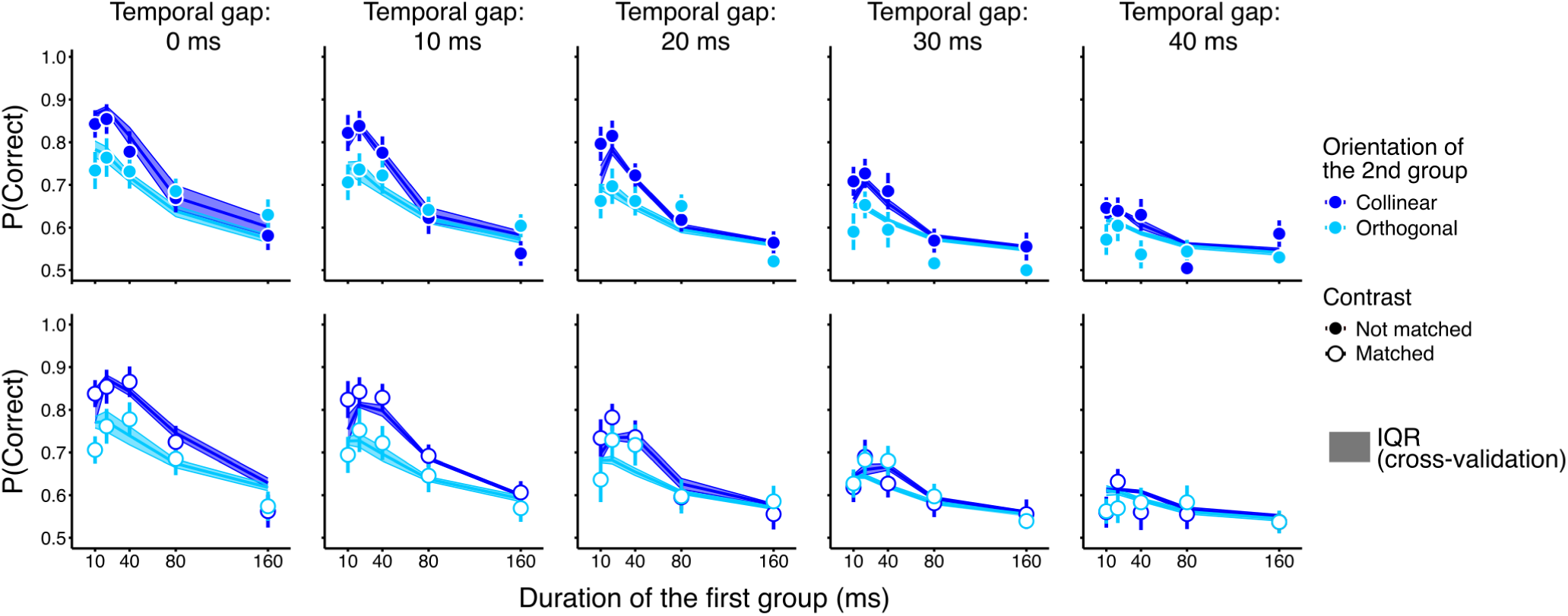
Model fits to data in Experiment 1. Fits of the Tuned Delayed Normalisation model are shown as lines, and shaded regions correspond to inter-quartile range (IQR) from jackknife cross-validation. Symbols show human performance (same as Fig.4). Performance for the five temporal gap conditions is shown in different columns, and the two contrast conditions in rows. Colour codes for the two relative orientation conditions (dark and light blue, orientation of the second group collinear or orthogonal, respectively).

**Table 1.** Model parameters for the two experiments. Values in bold indicate fixed parameters. Inter-quartile range of parameters is obtained by jackknife resampling method.

| Parameter | Description | Experiment 1 | IQR | Experiment 2 | IQR |
| --- | --- | --- | --- | --- | --- |
| $\tau_1$ | Time-constant for TIR | **0.03** | - | **0.03** | - |
| w | Weight on the delay of the negative lobe of TIR | **0.4** | - | **0.4** | - |
| $\tau_2$ | Exponential decay constant | $4.34 \cdot 10^{-2}$ | $4.1 - 5.5 \cdot 10^{-2}$ | $5.32 \cdot 10^{-2}$ | $5.2 - 5.44 \cdot 10^{-2}$ |
| $\tau_3$ | Exponential delay constant | **0.08** | - | **0.08** | - |
| m | Exponent | **1** | - | **1** | - |
| $\sigma$ | Semi-saturation constant | $2 \cdot 10^{-3}$ | $1.89 - 5.33 \cdot 10^{-3}$ | $3.7 \cdot 10^{-3}$ | $3.4 - 4.7 \cdot 10^{-3}$ |
| $\sigma_s$ | Spatial standard deviation | 1.12 | 1.10-1.25 | 13.9 | 11.7 - 14.9 |
| $\gamma_u$ | Weight of the untuned pool | 1.35 | 1.19-1.65 | 0.270 | 0.267-0.273 |
| $\gamma_c$ | Weight of tuned pool | **1** | - | **1** | - |
| p1 | Detection threshold | 0.427 | 0.149-0.352 | $8.95 \cdot 10^{-2}$ | $8.5 - 10.2 \cdot 10^{-2}$ |
| p2 | Detection slope | 293 | 173-300 | 292 | 282-299 |
| p3 | Decision threshold | 0.625 | 0.664-0.750 | 0.464 | 0.435-0.487 |
| p4 | Decision slope | 1.4740 | 1.52-1.87 | 0.989 | 0.948-1.04 |

### Temporal integration depends on how signals are delivered over time

We showed that inhibition over time and space can explain effects of duration and collinearity on temporal integration. From the time-course of the inhibition, we could predict the effects of the distribution of elements in the two groups across time. When more elements are presented in the first group, there is more inhibition earlier in time, relative to the peak of the drive. This might lead to better performance, since the effect of inhibition on the visibility will be smaller (the drive is stronger at the moment of the strongest inhibition). To test this prediction, we presented a new set of temporally segmented ring stimuli in a second experiment. In the first and second groups, the grating elements were oriented so as to form a collinear contour, or were randomly oriented, thus resulting in four experimental conditions: collinear-collinear, collinear-random, random-collinear, random-random (**Fig**.**6a**). Crucially, we also varied the proportion of elements presented in the first group (4, 8, 10, 12 or 16, out of 20 in total), with the complementary number in the second group (16, 12, 10, 8, or 4; **Fig**.**6b**). These five levels of element numbers were crossed with the four orientation conditions to allow us to test how the stability of the spatial structure affects temporal integration over time. To manage the duration of the experiment to a reasonable time per participant, we fixed the temporal parameters of the task (durations of the first and the second group to 20 ms and temporal gap to 10 ms, see Methods for details). The task was identical, namely to determine in a two-interval two-alternative forced choice which interval contained the stimulus that had a missing element.

Performance, and therefore temporal integration of the two groups, improved when the number of gratings presented in the first group increased, as we hypothesised (type III Wald *χ*^2^(2) = 20.646; **Fig**.**6c**, circles and error bars indicate human performance). The relative orientation of gratings in the display across the two groups affected temporal integration: performance was impaired when a small number of collinear elements in the first group were followed by a larger number of random elements in the second group (*χ*^2^(6) = 21.684, *p <* 0.01; **Fig**.**6c**, light blue symbols).

When the Tuned Delayed Normalisation model was fit to the human data, it accounted quantitatively for the two main features of the behaviour (**Fig**.**6c**, solid lines). In particular, there was an increase in performance with an increase in the number of gratings presented in the first group. The model also correctly reproduced the order of difficulty of the four spatial orientation conditions, with the collinear-random being the most difficult. Interestingly, performance for this latter condition was non-monotonic as a function of the number of elements in the first group, and the model also reproduced this feature. In Experiment 2, the nested hypothesis testing did not provide evidence that the full model was significantly better (*λ_Nested_* = −427.849, *λ_F ull_* = −425.844, *χ*^2^(3) = 4.01, *p* = 0.260). However, the nested model fails to account qualitatively for the pattern of data, since it predicts neither the effect of the proportion of elements presented in the first group nor the effect of the collinearity (see Supplementary Information).

### Model prediction across a variety of spatio-temporal contexts

In the final experiment, we examined how different expected performance can be for temporally segmented ring stimuli that simply very in the timing of the elements’ presentation. We simulated 10, 000 displays, where each display contained 16 gratings, lasted for an overall duration of 60 ms, and had a pseudo-random distribution of gratings over space and time. Gratings’ durations were selected randomly (from 10 ms to the remaining display time), and displays in which all gratings appeared simultaneously were excluded to avoid trivially easy trials (see Methods for more details). We used the best fit from Experiment 1 to predict performance for each of the displays, and selected the ten most difficult, the ten easiest, and a final group of ten displays with median difficulty (**Fig**.**7a**). We presented those selected displays to participants in a 2AFC task, and asked them to select the interval with the missing grating, as in previous experiments. Tests were generated by randomly choosing a location and removing the grating at that location for the duration of that trial. Displays were also randomly rotated from trial to trial to avoid learning effects. Average human performance aligned very well with the model predictions (*χ*^2^(2) = 312.1, *p <* 0.01; **Fig**.**7b**), and performance increased for the medium and easy displays relative to the performance on difficult displays (regression coefficients in Supplementary Information). This experiment non only validates our model for a variety of spatial and temporal contexts, but also offer a short battery of stimuli to further investigate the effects of spatial location and relative orientation in the future.

## Discussion

The work described here addresses the question of the spatio-temporal nature of temporal integration in the human visual system. Behavioural experiments revealed an intricate relationship between spatial and temporal features of the visual information. Integration of successive visual events is better when they are brief and appear close in time^7,20,52,53^, but also when they form a continuous contour. Importantly, these spatial effects are not symmetric in time – temporal integration is improved if collinear elements are presented in the second, rather than in the first part of the display. We show that temporal integration is sensitive to both the spatial properties and the number of visual events presented over time, suggesting a continuous rather than discrete process at early stages of visual processing. The results are well-described by a computational model that defines temporal integration as an overlap of signals evoked by individual visual objects. Crucially, these signals are outputs of a tuned divisive normalisation mechanism^10,23,25^, operating both in time and space, and integrated via coincidence detection ^52–54^. We further evaluated interactions between spatial and temporal context of the display by contrasting model predictions with human performance on novel displays with different spatio−temporal structure. In summary, we have presented here the spatio−temporal mechanism that underlies the temporal integration of early visual signals, and this spatio−temporal process ultimately determines the content of our perception.

### Tuned Delayed Normalisation

Across three experiments, human performance on the detectability of a missing element in temporally segmented ring stimuli was well accounted for by a model implementing tuned and delayed divisive normalisation computations^10,23,25^, coupled with coincidence detection^52–54^. The model predicts that spatio-temporal context affects the temporal integration, as the time-course of the response to each grating is modulated by activity of its neighbours. Stronger normalisation shortens duration or delays onset and peak of normalised responses, decreasing the probability of their detection and temporal overlap with responses to other gratings. These interactions over time account for the effects of duration of the first group and temporal gap between the two groups. It also accounts for previously documented effects of the duration of the second group of elements^52,74^ (see Supplementary Information).

In the visual cortex, normalisation could be implemented either by lateral connections^10–12^ or feedback from higher visual areas circuits^57,75^, although a feedforward normalisation has also been proposed^76^. While our model is agnostic to specific mechanisms, its biological plausibility is supported by evidence that visual responses in the human visual cortex are well described by normalisation models, at least at the level of the population activity^23,25,71,77^. These computations might be implemented by means of recurrent circuits, with coupled neural integrators implementing normalisation via recurrent amplification^78^.

The model distinguishes two different contributions to the normalisation pool, with different time-constants. Specifically, the improvement of performance for collinear displays is implemented by two parameters: different contribution of the activity of non-collinear gratings to the normalisation pool (*γ_u_* > *γ_c_*) and a faster normalisation by collinear elements (*τ*_2_ *> τ*_3_, see Table 1). These are likely combinations of signals originating from different feedback connections, with distinct tuning and time-courses at different levels of the processing hierarchy^6,12,51,57,79^. Time constant of the collinear contribution to the pool was found to be faster relative to the untuned contribution in both experiments. The fast normalisation for the collinear normalisation pool aligns with findings showing fast feedback from extra-striatal visual areas that are likely to contribute to coding of collinearity^57,75,80^, and the time course or figure-ground segregation in areas V4 and V1^35,79^.

The parameters controlling the spatial extent and relative magnitude of the contribution of the untuned component to the normalisation pool were different in the first two experiments. Specifically, spatial spread of the pool was greater in Experiment 2, and contribution of the untuned component smaller. The top-down influences can be task or context-dependent^6,30,81^, and surround suppression could be a flexible mechanism, promoting heterogeneity in the input^6,71^. Pooling information across a larger spatial scale might be needed to account for spatial interactions between sparsely distributed units in conditions when only 2 or 4 gratings are presented in the first or the second group in Experiment 2. A smaller contribution of untuned gratings in Experiment 2 (randomly orientated) could be a consequence of the processing strategy of promoting variance of the inputs in uncertain contexts^6^. These hypotheses could be tested in future work, and refinement of the model could systematically disentangle the temporal dynamics of different contributions to the normalisation pool, and quantify the contributions of distinct neural pathways, such as long-range horizontal connections and top-down feedbacks.

### Relation to previous work

The model predictions align with some previous findings. For example, spatial separation of individual elements in a display has been shown to affect their temporal integration, with an increasing separation leading to better temporal integration^82^. These results are qualitatively in agreement with predictions of the model, since greater separation of elements would lead to less inhibition, longer responses and increased probability of their temporal overlap. In the masking literature, temporal asymmetry in lateral interactions between a target and collinear context has been documented^42^. Lateral facilitation of target detection by collinear context occurs when the target is presented after, but not before the context, similarly to the results observed here in Experiment 2. Furthermore, sensitivity of explicit judgements of the simultaneity of two visual events is impaired when they are grouped with a static collinear context^83^. Our findings and model predictions are in general agreement with these findings, as the temporal integration window is prolonged for collinear elements, rendering explicit judgements of synchrony more difficult.

One previous extension of the divisive normalisation model could account for effects of contextual facilitation in a contrast matching task, in the form of a multiplicative enhancement of the drive component of the response^84^. However, the enhancement did not depend on the collinearity between the center and surround in the periphery (although collinearity is important for contrast detection at threshold^29,85–87)^, and for the high contrasts used in our stimuli, this kind of enhancement is not expected, suggesting the existence of a different mechanism than the one we observed. Recent work^88^ demonstrated that delayed normalisation can emerge in a recurrent network model of neural responses. The network contains excitatory and suppressive drives that depend on recent input, within temporal windows. This work is a valuable step forward in describing moment-to-moment dynamics of context-dependent responses in the visual system. The present work goes beyond, by establishing a link between the putative neural computations and content of human visual experience.

### Relation to other models of temporal integration

Several models have previously described human behaviour in visual temporal integration tasks. Temporal integration was modelled as a correlation of filtered responses to the successive displays^52,53^ or their relative overlap^89^. These models are conceptually similar to the model proposed here, and predict several features of the data, such as the decrease of temporal integration with duration of the displays or temporal gap. However, our formulation has several advantages. Responses of the visual system to the presented displays are predicted with a biologically inspired model^12^ involving neural computations with a strong support in recent human electrocorticographic data^23,25,77^. Furthermore, the signals are combined by means of a coincidence detection in a robust manner that is independent of features of the visual stimulus such as contrast or specific shape of temporal impulse response. The effects of several parameters, such as duration, contrast or spatial proximity on temporal integration have been predicted by a neural network with reverberatory activations persisting beyond stimulus offsets, affected by a balance of excitation and inhibition within a neural network^90,91^. While the proposed network architecture may account for early visual responses, our model offers a parsimonious explanation of various effects, such as the influence of both the first and second element durations, using a single mechanism that is easy to interpret, and providing a quantitative fit to the data^89^.

The model proposed here defines temporal integration as a continuous process over space and time. This approach challenges theories that propose discrete sampling of visual information within successive and non-overlapping temporal windows, and provides constraints on where in the cortical hierarchy these mechanisms might operate^18–20,92^.

### Future extensions of the framework

Visual information is integrated at different spatial scales across the visual field and at different levels in the processing hierarchy^93,94^. Here we have modelled the spatial extent of the normalisation pool, but future extensions of the framework could be easily added to include spatial receptive fields of different shapes or sizes, as one expects to find in the peripheral visual field. Furthermore, including noise at different stages in the model could offer further insights into the limits of temporal integration in the visual system. While the sensory component in the model is dynamic, the perceptual decision assumes for now an integration of all the evidence over time equally. However, in natural scenarios, it is plausible that relevant signals would be extended over time, and vary in their intensity or quality. Future refinements of the model should also address the important issue for the visual system to decide when to start and stop integrating the relevant information, and to introduce memory and selection mechanisms^95,96^. In this project, we focused on temporal integration, a process that reduces the effects of stimulus and neural noise, and increases the robustness of visual representations in the visual system. However, to identify when an event occurs or to establish the order of a sequence of events, the human visual system must separate information across time and space. This segmentation in space and time is a challenge for future extensions of the model presented here within a unified framework^50,97,98^.

In summary, we report novel behavioral evidence for some complex interactions between temporal and spatial features in the integration of visual information. We propose a unifying computational framework that accounts both for these novel effects as well as previously established effects of temporal features. A substantial body of work has described the effects of spatial context in spatial integration in animal neurophysiology and human behaviour, both of which have been successfully modelled within the divisive normalisation framework^9,12,37,57,64,68^. Here, we build upon this seminal line of research by extending it to temporal processing, further narrowing the gap between the brain activity and human perception and behaviour^99^. Divisive normalisation has functional roles, in particular as an optimal computational strategy for decorrelating inputs present in natural images^2,62,100,101^, and increasing the stability of neural firing^102^. Our findings show how these computations affect information integration over time, segregating spatially adjacent signals, and promoting temporal integration of signals to contribute to the perception of an object.

## Methods

### Participants

Twenty participants were recruited for Experiment 1 (mean age 29.1 (sd = 4.4) years). One participant did not complete the two sessions, and data from another participant was excluded because they did not comply with the instructions (their performance was at chance even for the easiest conditions). In total, the performance of 18 participants was analysed. In Experiment 2, 18 participants participated (mean age 29.7 (sd = 3.98) years). Data from three participants was not included in the analyses, since their performance was at chance (*>* 60% correct) for at least one of the conditions tested. In Experiment 3, nine participants performed the task (mean age 23.6 (sd = 4.8)). All participants had normal or corrected-to-normal vision. The experiment was conducted in agreement with the Declaration of Helsinki and local ethics regulations, and all participants gave their written consent.

### Materials

Stimuli were presented on a CRT Sony Triniton (800 x 600 pix, refresh rate 100 Hz). The viewing distance was 60 cm and a chin-rest was used to minimise head movements. Experiments were created using Matlab R2024 and Psychtoolbox-3^103–105^ running on a MAC Mini. The analysis was conducted in R Studio environment^106^, with lme4 package^107^. Experiments were conducted in a dark room.

### Stimuli

The stimuli consisted of 2D sinusoidal gratings, size 1 degree of visual angle (dva), with spatial frequency 2 cycles per degree (cpd). Gratings were arranged on a virtual circle with 5 dva radius, centered at the center of the screen. A white fixation dot (0.5 dva) was presented at the center of the screen during the trial. They were presented on a mid-gray background (∼ 30 cd*/*m^2^). Sixteen gratings were presented in Experiments 1 and 3, and 20 in Experiment 2.

### Procedure and task

#### Experiment 1

On each trial, two arrays of sinusoidal gratings were presented on a virtual circle centered on the fixation point in two sequential intervals. In each interval, the gratings were randomly split in two equally sized groups, presented sequentially (**Fig 1a**). The test array was created by omitting one grating in one of the two intervals. Participants were asked to choose the interval that contained the test array, in a two interval two alternative forced-choice paradigm (2AFC). We varied the duration of the first group (10, 20, 40, 80 and 160 ms) and the temporal gap between the two groups (0, 10, 20, 30, 40 ms). The duration of the second group was always 10 ms (**Fig 1d**).

The orientation of the gratings’ carriers was either tangential to the virtual circle in both groups, forming a collinear contour, or the gratings’ orientations were collinear to the virtual circle in the first, and orthogonal in the second group (forming a disjoined contour, **Fig 1b**). All gratings had the same phase, that was randomly chosen on each trial. In one session the contrast of the first group was the same as the contrast of the second group (100%), and in the second session the contrast was reduced to match the perceived contrast of the second group (**Fig 1c**). Values were individually chosen based on a calibration session. The four conditions were presented in two sessions (on two separate days). The two spatial conditions (disjoined or continuous contours) were tested in separate blocks. In total, participants completed 2400 trials (5 durations of the first group x 5 temporal gaps x two spatial conditions x two contrast conditions x 24 repetitions), in two 1h sessions tested on different days.

In the calibration session, participants were presented with two successive rings of 16 collinearly arranged gratings. Durations of the two arrays corresponded to durations of the first and second group of gratings in the main experiment (the first array 10−160 ms and the second fixed to 10 ms). Contrast of the second array was fixed (100%), and the contrast of the first was adjusted based on participants’ responses (Accelerated Stochastic Approximation^108^) to match the perceived contrast of the second array. Staircases of two participants did not converge, and for those participants we used average matched contrast of other participants.

#### Experiment 2

The procedure was similar to that in Experiment 1, with several differences. Durations of the two groups were fixed to 20 ms and temporal gap to 10 ms. We varied the relative orientation of the gratings in four blocks. There were two possible arrangements: the gratings were either collinear to the virtual circle or randomly oriented (orientation of each grating randomly sampled from the range 20 − 160 degrees away from the collinear). In four different blocks, the collinear or random orientation could be presented in the first or in the second group (collinear-random, random-collinear, collinear-collinear, random-random, **Fig 6a**). Furthermore, in five different conditions in each block, we varied the number of gratings presented in the first group (4, 8, 10, 12 or 16). The blocks were completed in a counterbalanced order. Participants completed 48 trials for each of the conditions (4 relative orientation combinations x 5 group sizes x 48 repetitions), yielding 960 trials. The whole experiment lasted approximately one hour.

**Figure 6.**
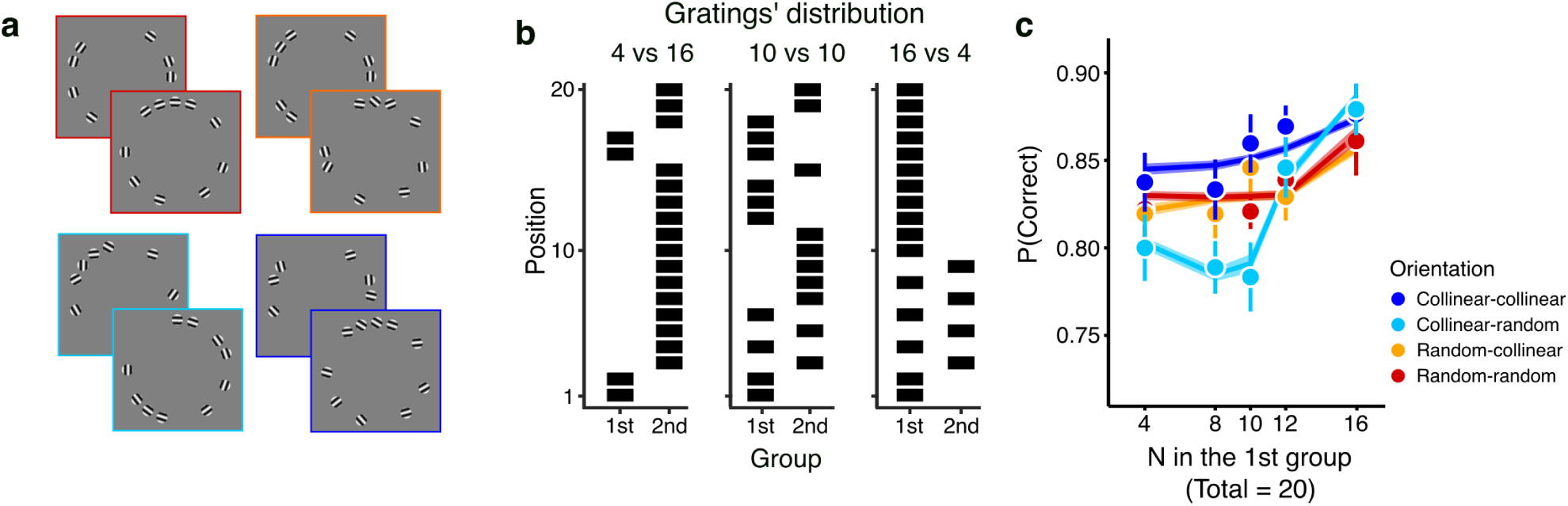
Stimuli and results of Experiment 2. a. To test the critical effects of spatial position and relative orientation on temporal integration, we varied the number and the nature of the elements in the two groups. The gratings were arranged to either form a collinear contour, or were randomly oriented, both in the first and in the second group, so as to create four conditions (color-coded): collinear-collinear, collinear-random, random-collinear, and random-random. b. The number of gratings presented in the first and the second group could take one of five levels (4, 8, 10, 12 or 16 out of 20 gratings in total in the first group, and the complementary number in the second group). The three panels illustrate the temporal distributions of the gratings across the two groups for three of these five levels (in which 4, 10 and 16 gratings were presented in the first group). c. Average percent correct is plotted as a function of the first group’s size, separately for the four spatial conditions (color coded). Temporal integration improved as the number of gratings presented in the first group increased. It was impaired when collinear elements were followed by random elements in the second group (light blue symbols). Symbols show human performance (error bars are standard error of the mean). Lines show the model fits, and shaded regions correspond to IQR from jackknife cross-validation.

#### Experiment 3

In Experiment 3, the procedure was similar to that in Experiment 1, with several modifications. Instead of randomly choosing gratings’ locations as in previous experiments, in Experiment 3 each participant performed the temporal integration task on the same 30 stimuli. In order to choose stimuli displays we first randomly generated 10, 000 displays. For each display, 16 gratings were presented and the display duration was fixed to 60 ms. Every 10 ms, the gratings to be displayed were chosen randomly. Their duration was also randomly sampled (from 10 ms to the remainder of the display duration, in 10 ms steps), and carrier orientation was tangential to the outline of the ring. In order to eliminate trivially easy displays, we excluded stimulated displays in which all gratings were presented at the same time. The parameters that best fit average data of Experiment 1 were used to predict performance on the simulated displays. Then, we selected 30 displays: the most difficult (10), the easiest (10) and the displays of median difficulty (10) as stimuli in Experiment 3 (see **Fig 7a** for examples). We did not constrain total energy across space and time, but the performance predicted by the model did not depend on this variable (**Fig S4**). In the experiments, on each presentation the stimulus display was randomly rotated in order to minimise learning. In total, participants completed 720 trials in ∼ 30 minutes.

**Figure 7.**
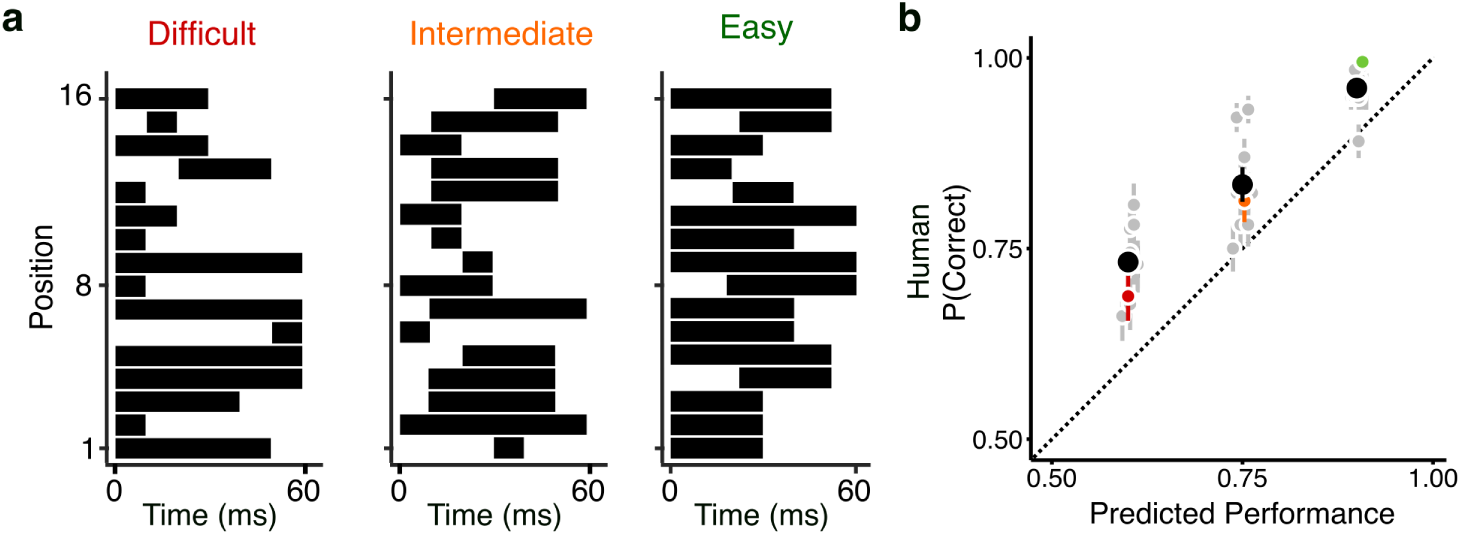
Stimuli and results of Experiment 3. **a**. Examples of spatial and temporal distributions of gratings in displays presented in Experiment 3. The three panels show one example of a difficult display (left), a display with medium difficulty (middle) and one easy (right). **b**. Average human performance in Experiment 3 is shown against model predictions. The performance difficulty on the novel displays was correctly predicted by the model using the parameters obtained in Experiment 1. Black symbols show performance averaged across participants and displays, and gray symbols show performance for individual displays (10 in each of the three chosen difficulty levels). Error bars indicate standard error of the mean. The red, orange and green symbols show performance on the three displays illustrated in panel **a**.

### Data analysis

We analysed participants’ responses in the three experiments by means of a generalised linear mixed-effect models (binomial regression with logit link). Significance of the predictors was assessed by means of Wald’s *χ*^2^ test (type III), and *β* coefficients are reported as the effect size measure. In Experiment 1, fixed effects of the four predictors and their interactions were tested. The random structure consisted of random intercept and slopes for the contrast of the first group and orientation of the second group at the level of participant were included, to account for additional variability. Given the non-linear relationship between the duration of the first group and the performance (**Fig. 2**), we modelled the effect of the duration of the first group as a second order polynomial. In Experiment 2, we tested fixed effects of the relative orientation and the proportion of gratings in the first group (included as a second order polynomial) and their interaction on the performance. We included a random intercept and slope for the relative orientation condition at participant level. In Experiment 3 we tested the effect of the predicted difficulty level (categorical fixed effect) on the performance and included random intercept at the level of participant.

### Model

#### Simulation procedures

To obtain model prediction for each of the conditions, trials were simulated for each condition (24 or 48 in Experiments 1 and 2, same as the number of trials in the experiment). Each simulated trial lasted 1.5 s with time steps Δ t of 1 ms. For each trial and each condition, displays were generated by randomly assigning each of the gratings to the first or the second group (half in Experiment 1 or 0.2, 0.4, 0.5, 0.6, 0.8 proportion in Experiment 2). For each simulated trial, responses to each grating were predicted by the Tuned Delayed Normalisation model, and then their temporal overlap was calculated as a product of the predicted time-courses (see eq. 10). Evidence for temporal overlap was integrated over time, averaged for all the trials, and passed through a logistic function to obtain predicted percent of correct responses for that condition. The model simulations and parameter estimation were run using MATLAB.

#### Fitting procedures

The model was fit to the average data (percent correct, 100 data points for Experiment 1 and 20 for Experiment 2) by minimising the negative log-likelihood of the model given the data with *fminsearchbnd*. The set of parameters that minimised the cost function was chosen as the best fit. Table 1 shows parameters that best fit the behavioural data. Parameters that were fixed by hand are shown in bold, and do not have confidence intervals. Values of these parameters were chosen based on reported values based on model fits to electrocorticographic recordings of human V1^25^. Confidence intervals were obtained by jackknife resampling procedure^109^, a conservative resampling method suitable for small sample sizes. Performance of one participant was excluded at the time, and fitting procedure repeated on these resampled averages (18 repetitions for Experiment 1 and 15 for Experiment 2).

To evaluate model performance, we used a likelihood ratio test for nested models, *λ_LR_* = –2(*λ_Nested_* − *λ_F ull_*), where *λ_Nested_* and *λ_F ull_* correspond to log likelihoods of the models to be compared. *λ_LR_* follows a *χ*^2^ distribution with degrees of freedom equal to the difference in the number of parameters between the two models. Small p-values indicate that the more complex model provides significantly better fit to the data.

