## Supplemental Information for "Spatial and temporal contexts drive visual temporal integration via spatiotemporal normalisation"

Laboratoire des systèmes perceptifs, Département d'études cognitives, École normale supérieure, PSL University, CNRS, Paris, France

### 1. Experiment 1

#### 1.1. Generalised linear mixed-effect model

To quantify results in Experiment 1, we submitted the data to a generalised linear mixed-effect model, and tested effects of the duration of the first group, ISI, contrast and relative orientation condition and their interactions on the participants' ability to detect the interval with the missing grating (binomial regression with logit link). The random structure consisted of random intercept and slopes for the contrast of the first group and orientation of the second group at the level of participant, to account for additional variability. Given the non-linear relationship between the duration of the first group and the performance (see Fig S2), we modelled the effect of the duration of the first group as a second order polynomial (Duration and Duration$^2$ in Table S1). Coefficients for all the fixed effect tested are shown in Table S1.

Duration of the inter-stimulus interval negatively affected temporal integration (type III Wald $\chi^2(1) = 1650$, $p < 0.01$, $b_{ISI} = -44.36$, SE = 1.092, $z = -30.27$, $p < 0.01$). More surprisingly, but in line with previous work[7], the performance non-linearly decreased with the duration of the first group ($\chi^2(2) = 1077$, $p<0.01$, linear effect: $b_{Linear} = -14.66$, SE = 0.485, $z = -30.27$, $p < 0.01$, quadratic: $b_{quadratic} = 19.7$, SE = 1.855, $z = 10.62$, $p < 0.01$). There was also a significant interaction between the duration of the first group and the ISI ($\chi^2(2) = 13072$, $p<0.01$) indicating that the effect of the duration was smaller at longer ISIs ($b_{ISI\ x\ Linear} = 266.510$, SE = 2.432, $z = 109.5$, $p < 0.01$, $b_{ISI\ x\ Quadratic} = -85.36$, SE = 1.804, $z = -47.273$, $p < 0.01$).

Importantly, the spatial orientation of the second group also had an effect ($\chi^2(1) = 273.52$, $p<0.01$). The performance was better when both groups had collinearly arranged elements ($b_{orthogonal} = -0.973$, SE = 0.063 $z = -15.412$, $p < 0.01$). The collinearity interacted with the duration of the first group (interaction effect, $\chi^2(2) = 405.83$, $p<0.01$) and ISI (interaction effect, $\chi^2(1) = 273.8$, $p<0.01$). The effect of the duration of the first group (linear effect: $b_{Linear\ x\ Orthogonal} = 12.61$, SE = 0.654, $z = 19.25$, $p < 0.01$, quadratic: $b_{Quadratic\ x\ Orthogonal} = -41.24$, SE = 2.416, $z = -17.076$, $p < 0.01$) and ISI ($b_{ISI\ x\ Orthogonal} = 23.72$, SE = 1.436, $z = 16.52$, $p < 0.01$) were smaller when the second group contained orthogonal elements.

We found no evidence for an effect of the contrast of the first group (contrast identical to that of the second group or perceptually matched; $\chi^2(1) = 0.91$, p=0.34, $b_{contrast} = 0.063$, SE = 0.065). However, there were significant interactions between the contrast of the first group and its duration ($\chi^2(2) = 759.34$, p<0.01), as well as ISI ($\chi^2(1) = 79.91$, p<0.01). These effects suggest that when the contrast of stimuli was not adjusted, the performance declined faster with duration of the first group (linear effect: $b_{Linear\ x\ Not\text{-}mached} = -9.640$, SE = 0.600, z = -16.086, p < 0.01, quadratic: $b_{Quadratic\ x\ Not\text{-}mached} = 54.57$, SE = 1.984, z = 27.504, p < 0.01), and longer ISIs ($b_{ISI\ x\ Not\text{-}mached} = 10.840$, SE = 1.212, z = 8.939, p < 0.01).

There was a complex relationship between the tested variables reflected in higher-order interactions. We found evidence for a significant three-way interaction between the duration of the first group, ISI and orientation of the second group ($\chi^2(2) = 33015.06$, p<0.01, linear: $b_{Linear\ x\ ISI\ x\ Orthogonal} = -194.3$, SE = 2.450, z = -79.34, p < 0.01, quadratic: $b_{Quadratic\ x\ ISI\ x\ Orthogonal} = 283.8$, SE = 1.881, z = 150.85, p < 0.01), indicating that the interaction between the duration of the first group and the ISI was smaller in the condition where the second group had orthogonal elements.

There was also an interaction between the duration of the first group, ISI and contrast of the first group ($\chi^2(2) = 7792$, p<0.01, linear: $b_{Linear\ x\ ISI\ x\ Not\text{-}matched} = -8.200$, SE = 1.734, z = -4.728, p < 0.01, quadratic: $b_{Quadratic\ x\ ISI\ x\ Not\text{-}matched} = -188.7$, SE = 2.134, z = -88.21, p < 0.01), suggesting that the interaction between the duration of the first group and the ISI was smaller in the condition where the contrast of the first group was not perceptually matched.

A significant three-way interaction between the duration and the contrast of the first group and the orientation of the second group ($\chi^2(2) = 442.6$, p<0.01, linear: $b_{Linear\ x\ Orthogonal\ x\ Not\text{-}matched} = 7.810$, SE = 0.747, z = 10.45, p < 0.01, quadratic: $b_{Quadratic\ x\ Orthogonal\ x\ Not\text{-}matched} = -37.31$, SE = 1.782, z = -20.94, p < 0.01), indicated that the decrease in the effect of the duration of the first group in the condition in which the second group contained orthogonal elements was further decreased for not-matched stimuli. In contrast, a significant three-way interaction between the ISI and the orientation of the second and the contrast of the first group ($\chi^2(1) = 58.45$, p<0.01, $b_{ISI\ x\ Orthogonal\ x\ Not\text{-}matched} = -10$, SE = 1.394, z = -7.645, p < 0.01) indicated that the decrease in the effect of ISI for stimuli containing orthogonal elements was greater for matched stimuli. Finally, a significant four-way interaction between all the variables tested $\chi^2(2) = 61931$, p<0.01, $b_{Linear\ x\ ISI\ x\ Orthogonal\ x\ Not\text{-}matched} = -83.09$, SE = 2.144, z = -38.761, p < 0.01, quadratic: $b_{Quadratic\ x\ ISI\ x\ Orthogonal\ x\ Not\text{-}matched} = 420$, SE = 1.709, z = 245.8, p < 0.01), suggests that the negative effect of orthogonal orientation of the second group on interaction between the duration of the first group and ISI was stronger when stimuli were not matched (contrast was higher).

**Table S1. Coefficients of the generalised linear mixed-effect model in Experiment 1.**

| | $\beta(SE)$ |
|---|---|
| (Intercept) | $2.171^{***}$ |
| | (0.097) |
| Duration | $-14.661^{***}$ |
| | (0.484) |
| Duration$^2$ | $19.707^{***}$ |
| | (0.419) |
| ISI | $-44.36^{***}$ |
| | (1.09) |
| Orientation (coll-orth) | $-0.97^{***}$ |
| | (0.06) |
| Contrast (not-matched) | $0.0632^{*}$ |
| | (0.065) |
| Duration:ISI | $266.1^{***}$ |
| | (2.42) |
| Duration$^2$ :ISI | $-85.361^{***}$ |
| | (1.8) |
| Duration:Orientation (coll-orth) | $12.61^{***}$ |
| | 0.65) |
| Duration$^2$ :Orientation (coll-orth) | $-41.24^{***}$ |
| | (2.41) |
| ISI:Orientation (coll-orth) | $23.71^{***}$ |
| | (1.43) |
| Duration:Contrast (not-matched) | $-9.64^{***}$ |
| | (0.60) |
| Duration$^2$ :Contrast (not-matched) | $54.57^{***}$ |
| | (1.98) |
| ISI:Contrast (not-matched) | $10.83^{***}$ |
| | (1.21) |
| Orientation (coll-orth) :Contrast (not-matched) | $-0.041$ |
| | (0.07) |
| Duration:ISI:Orientation (coll-orth) | $-194.25^{***}$ |
| | (2.45) |
| Duration$^2$:ISI:Orientation (coll-orth) | $283.811^{***}$ |
| | (1.88) |
| Duration:ISI:Contrast (not-matched) | $-8.19^{**}$ |
| | (1.733) |
| Duration$^2$:ISI:Contrast (not-matched) | $-188.656^{***}$ |
| | (2.138) |
| Duration:Orientation (coll-orth):Contrast (not-matched) | $7.81^{***}$ |
| | (0.75) |
| Duration$^2$:Orientation (coll-orth):Contrast (not-matched) | $-37.31^{***}$ |
| | (1.78) |
| ISI:Orientation (coll-orth):Contrast (not-matched) | $-10.65^{***}$ |
| | (1.39) |
| Duration:ISI:Orientation (coll-orth):Contrast (not-matched) | $-83.08^{***}$ |
| | (2.14) |
| Duration$^2$:ISI:Orientation (coll-orth):Contrast (not-matched) | $419.98^{***}$ |
| | (1.70) |
| AIC | 52754.11 |
| BIC | 53014.32 |
| Log Likelihood | $-26347.06$ |
| Num. obs. | 43200 |
| Num. groups: Participant | 18 |
| Var: Participant (Intercept) | 0.14 |
| Var: Participant Orientation (coll-orth) | 0.01 |
| Var: Participant Contrast (not-matched) | 0.02 |
| Cov: Participant (Intercept) Orientation (coll-orth) | $-0.01$ |
| Cov: Participant (Intercept) Contrast (not-matched) | $-0.02$ |
| Cov: ParticipantOrientation (coll-orth) Contrast (not-matched) | 0.00 |

$^{***}p < 0.001$; $^{**}p < 0.01$; $^{*}p < 0.05$

### 1.2. Temporal integration with untuned Delayed Normalisation

To test whether additional complexity introduced to the Delayed Normalisation Model was justified by the data, we and performed a nested hypothesis test. We tested performance of a simple model, where we the structure of the decisional part of the model (detection and integration) identical, and modified the normalisation mechanism, following the model structure reported in previous work[1]. Pool for the response to a grating $i$ is a rectified and exponentiated linear response to the grating $L_i$ convolved with the exponential decay function $h_2$.

$$Pool_i = |L_i * h_2(\tau_2)|^m \qquad \text{(Equation 1)}$$

A nested hypothesis test (likelihood ratio test) indicated that the full model described the data significantly better ($\lambda_{Nested}$ = -1496.51, $\lambda_{Full}$ = -1490.8, $\chi^2(3)$= 11.42, $p < 0.01$). Qualitatively, the model predicts effects of the duration and ISI, but fails to account for differences in the relative orientation of between the gratings across groups and contrast (Fig S1). Parameters that provided the best fit to data are shown in Table S2.

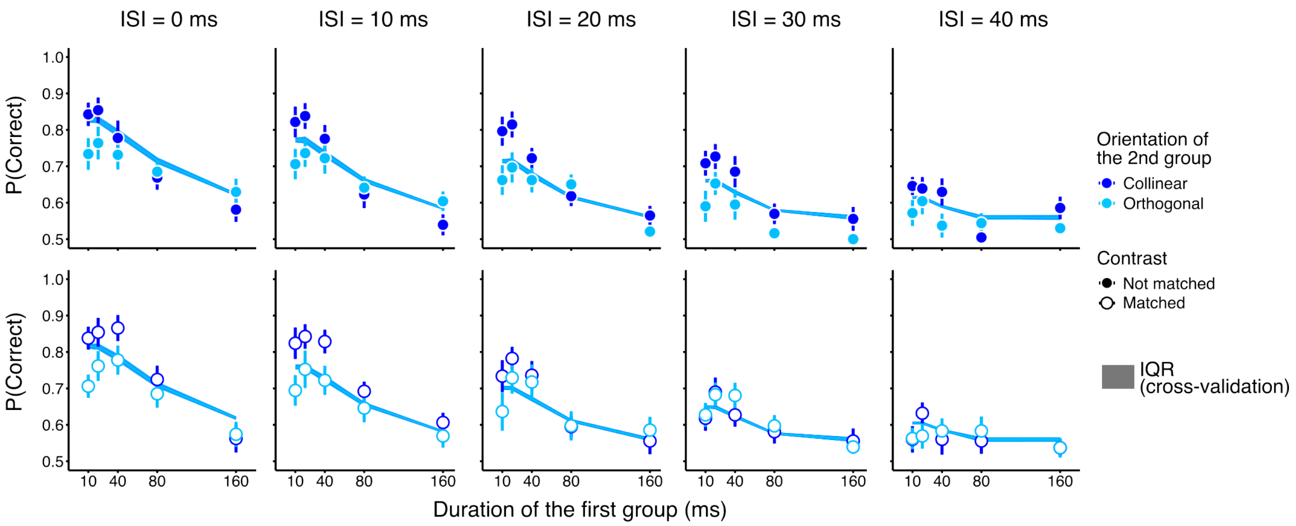


**Fig S1. Fits of the Delayed Normalisation model to data in Experiment 1.** Fits of the Delayed Normalisation model are shown in lines, and shaded regions correspond to IQR from jackknife cross-validation. Data are same as shown in Fig 4 and 5 in the manuscript. Performance for the five ISI conditions is shown in columns, and the two contrast conditions in rows. Colour codes for the two relative orientation conditions (dark and light blue, orientation of the second group collinear or orthogonal, respectively).

**Table S2. Parameters of the (untuned) Delayed Normalisation model providing best fit to the data in Experiment 1.**

| **Parameter** | $\tau_1$ | w | $\tau_3$ | m | $\sigma_n$ | $p_1$ | $p_2$ | $p_3$ | $p_4$ |
|---|---|---|---|---|---|---|---|---|---|
| **Value** | **0.03** | **0.4** | **0.08** | **1** | 0.01 | 0.937 | 299 | 0.751 | 1.14 |

#### 1.3. Supplementary Experiment 1: Effect of the duration of the second group

Previous work showed that when stimuli are presented in two groups sequentially (as in Experiments 1 and 2), temporal integration declines with an increase in the durations of both first and the second group[2,3]. To test whether the model can predict variations in performance with variation in the duration of the second group, we collected an independent dataset (N=5), and fit the model to the data. We presented 16 gratings sequentially in two groups, and all gratings formed a collinear contour. Inter-stimulus interval was always 0, and duration of both the first and the second group were varied in 5 steps (10, 20, 40, 80 and 160 ms) yielding 25 conditions. There were 24 repetitions for each of the condition. As expected[2,3], the duration of both the first and the second group affected temporal integration. The performance decreased with duration of either the first or the second group (Fig S2). Next, we fitted the data with the Tuned Delayed Normalisation model of temporal integration. The model successfully predicted the performance decrease with the increase in the duration of both the first and the second group (Fig S2, lines), although it slightly underestimates performance when duration of the second group is 160 ms. Parameters that provided the best fit to data are shown in Table S3.

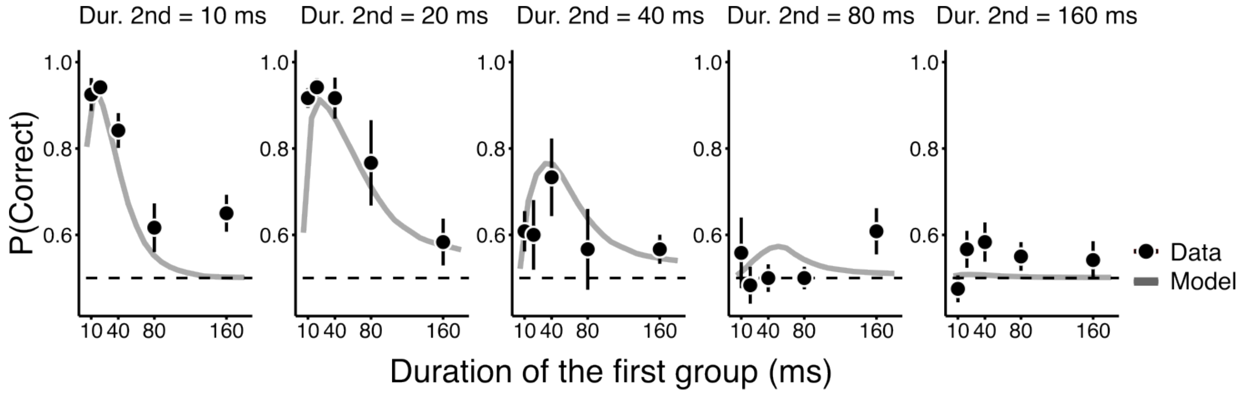


**Fig. S2. Results of Experiment S1 and model fit.** Each panel shows the average percent correct against the duration of the first group (black symbols). Performance for the five durations of the second group are shown in different panels. Performance decreased as a function of the durations of both the first and second groups. Fits of the Delayed Normalisation model are shown in gray lines lines. Error bars show standard error of the mean.

**Table S3. Parameters providing best fit to the data of Supplementary Experiment 1.**

| **Parameter** | $\tau_1$ | w | $\tau_2$ | $\tau_3$ | m | $\sigma_n$ | $\sigma_s$ | $\gamma_u$ | $\gamma_c$ | $p_1$ | $p_2$ | $p_3$ | $p_4$ |
|---|---|---|---|---|---|---|---|---|---|---|---|---|---|
| **Value** | **0.03** | **0.4** | $3.9\ 10^{-2}$ | **0.08** | **1** | $2\ 10^{-4}$ | 1.014 | **-** | **1** | 0.469 | 299 | 0.78 | 3.99 |

### 2. Experiment 2

#### 2.1. Generalised linear mixed-effect model equation and coefficients

In Experiment 2, the temporal integration of the two groups improved with an increase in the number of gratings presented in the first group. There was also an effect of the relative orientation of gratings in the display, and performance was impaired when collinear elements in the first group were followed by random elements in the second group. We tested effects of the relative orientation and the proportion of gratings in the first group (second order polynomial) and their interaction, with random intercept at the participant level. There was a significant effect of the proportion of stimuli in the first group (type III Wald $\chi^2$ (2) = 20.646, $p < 0.01$, linear: $b_{Linear}$ = 15.28, SE = 3.364, z = 4.542, $p < 0.01$, quadratic: $b_{Quadratic}$ = 1.757, SE = 3.737, z = 0.470, p = 0.638). There was evidence for a significant main effect of the relative orientation ($\chi^2(3)$ = 3.596, p=0.310). However, there was a significant interaction, indicating that when gratings were collinear in the first and random in the second group, the slope was steeper and more non-linear ($\chi^2(6)$ = 21.68, $p < 0.01$, linear: $b_{Linear}$: coll-rand = 11.90, SE = 3.702, z = 3.215, $p < 0.01$, quadratic: $b_{Quadratic}$: coll-rand = 15.14, SE = 4.306, z = 3.516, $p < 0.01$). Coefficients for all the fixed effect tested are shown in Table S4.

**Table S4. Coefficients of the generalised linear mixed-effect model in Experiment 2.**

| | $\beta(SE)$ |
|---|---|
| (Intercept) | $1.84^{***}$ |
| | (0.12) |
| Proportion | $15.28^{***}$ |
| | (3.36) |
| Proportion$^2$ | 1.76 |
| | (3.74) |
| Condition (coll-rand) | $-0.23$ |
| | (0.16) |
| Condition (rand-coll) | $-0.18$ |
| | (0.09) |
| Condition (rand-rand) | $-0.16$ |
| | (0.11) |
| Proportion:Condition (coll-rand) | $11.90^{**}$ |
| | (3.70) |
| Proportion$^2$:Condition (coll-rand) | $15.14^{***}$ |
| | (4.31) |
| Proportion:Condition (rand-coll) | $-3.12$ |
| | (7.71) |
| Proportion$^2$:Condition (rand-coll) | 1.84 |
| | (6.82) |
| poly(prop_mod, 2)1:Condition (rand-coll) | $-4.07$ |
| | (4.58) |
| Proportion$^2$:Condition (rand-rand) | 3.68 |
| | (7.58) |
| AIC | 12599.30 |
| BIC | 12765.95 |
| Log Likelihood | $-6277.65$ |
| Num. obs. | 14400 |
| Num. groups: Participant | 15 |
| Var: Participant (Intercept) | 0.17 |
| Var: Participant Condition (coll-rand) | 0.30 |
| Var: Participant Condition (rand-coll) | 0.06 |
| Var: Participant Condition (rand-rand) | 0.11 |
| Cov: Participant (Intercept) Condition (coll-rand) | $-0.11$ |
| Cov: Participant (Intercept)Condition (rand-coll) | $-0.05$ |
| Cov: Participant (Intercept) Condition (rand-rand) | $-0.05$ |
| Cov: Participant Condition (coll-rand) Condition (rand-coll) | 0.11 |
| Cov: Participant Condition (coll-rand)Condition (rand-rand) | 0.16 |
| Cov: Participant Condition (rand-coll) Condition (rand-rand) | 0.08 |

$^{***}p < 0.001$; $^{**}p < 0.01$; $^{*}p < 0.05$

### 2.2. Temporal integration with (untuned) Delayed Normalisation

As for Experiment 1, performed a nested hypothesis test, where the full model was compared to a simpler model (the simplified normalisation mechanism, following the model structure reported in previous work[1]). In Experiment 2, the nested hypothesis test did not provide evidence for the full model $\lambda_{Nested}$ = -427.8, $\lambda_{Full}$ = -425.8, $\chi^2(3)$= 4.01, p = 0.260). Nevertheless, this model fails to account qualitatively for the pattern of data, since it predicts neither the effect of the proportion of stimuli presented in the first group nor the effect of the collinearity (Fig S3). Parameters that provided the best fit to data are shown in Table S5.

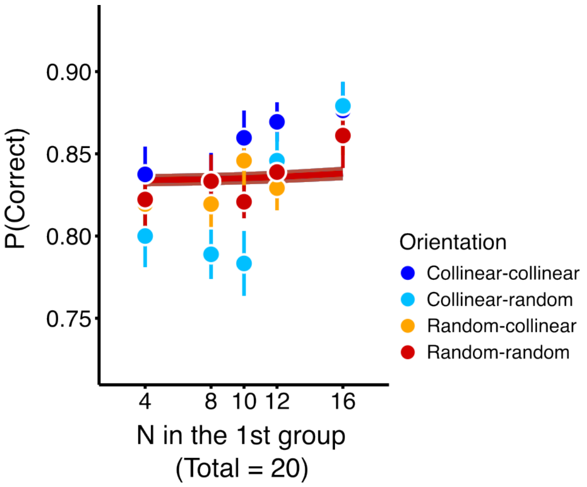


**Fig S3. Fits of the Delayed Normalisation model to data in Experiment 2.** Average percent correct (same data as in Fig 6c in the manuscript) as a function of the first group's size, separately for the four spatial conditions (color coded). Lines show the model fits, and shaded regions correspond to IQR from jackknife cross-validation. The model fails to capture effects observed in the data.

**Table S5. Parameters of the (untuned) Delayed Normalisation model providing best fit to the data in Experiment 2.**

| **Parameter** | $\tau_1$ | w | $\tau_3$ | m | $\sigma_n$ | $p_1$ | $p_2$ | $p_3$ | $p_4$ |
|---|---|---|---|---|---|---|---|---|---|
| **Value** | **0.03** | **0.4** | **0.08** | **1** | $9\ 10^{-4}$ | 0. 924 | 241 | 0.828 | 1.81 |

### 3. Experiment 3

#### 3.1. Generalised linear mixed-effect model equation and coefficients

We simulated 10,000 displays, where each display contained 16 gratings, lasted for 60 ms and had a pseudo-random distribution of gratings over space and time. Gratings' durations were selected randomly (from 10 ms to the remaining display time), and displays in which all gratings appeared simultaneously were excluded to avoid trivially easy trials (see Methods for more details). We used the best fit from Experiment 1 to predict performance on each of the displays, and selected the ten most difficult, the easiest and displays with median difficulty (Fig. 7a). We presented those selected displays to participants in a 2AFC task, and asked them to select the interval with the missing grating, as in previous experiments. Tests were generated by randomly choosing a location where no grating was presented on that trial, and displays were rotated from trial to trial to avoid learning effects. To quantify the results, differences in probability of correct responses as a function of the predicted difficulty were tested by means of a generalised mixed-effect (random intercept at the level of participant). Log odds of correct responses were different for the three categories of displays ($\chi^2(2) = 312.1$, $p < 0.01$), increasing for the medium ($b_{medium} = 0.633$, $SE = 0.082$, $z = 7.746$, $p < 0.01$) and easy displays ($b_{easy} = 2.24$, $SE = 0.129$, $z = 17.37$, $p < 0.01$) relative to the performance on difficult displays.

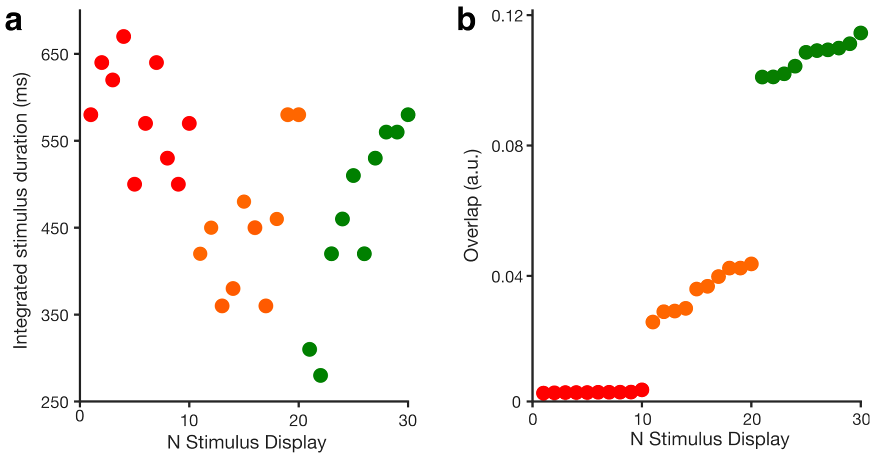


**Fig S4. Integrated contrast across displays used in Experiment 3. a.** Integrated contrast across time and the 16 locations is shown for the 30 stimuli used in Experiment 3. Color codes the difficulty category (red – high, orange – medium, green – low difficulty). **b.** Predicted overlap of the signals by the Tuned Delayed Normalisation model across the 16 locations.
